# Predicting primer and panel off-target rate in QIAseq targeted DNA panels using convolutional neural networks

**DOI:** 10.1101/2020.07.13.201558

**Authors:** Chang Xu, Raghavendra Padmanabhan, Frank Reinecke, John DiCarlo, Yexun Wang

## Abstract

In QIAseq targeted DNA panels, synthetic primers (short single-strand DNA sequences) are used for target enrichment via complementary DNA binding. Off-target priming could occur in this process when a primer binds to some loci where the DNA sequences are identical or very similar to the target template. These off-target DNA segments go through the rest of the workflow, wasting sequencing resources in unwanted regions. Off-target cannot be avoided if some segments of the target region are repetitive throughout the genome, nor can it be quantified until after sequencing. But if off-target rates can be prospectively predicted, scientists can make informed decisions about investment on high off-target panels.

We developed `pordle` (<u>p</u>redicting <u>o</u>ff-target rate with <u>d</u>eep learning and <u>e</u>pcr07), a convolutional neural network (CNN) model to predict off-target binding events of a given primer. The neural network was trained using 10 QIAseq DNA panels with 29,274 unique primers and then tested on an independent QIAseq panel with 7,576 primers. The model predicted a 10.5% off-target rate for the test panel, a -0.1% bias from the true value of 10.6%. The model successfully selected the better primer (in terms of off-target rate) for 89.2% of 3,835 pairs of close-by primers in the test panel whose off-target rates differ by at least 10%. The order-preserving property may help panel developers select the optimal primer from a group of candidates, which is a common task in panel design.

## 1. Introduction

The QIAseq targeted DNA panels utilize a PCR-based enrichment workflow to turn amplicons that represent regions of interest into library fragments for sequencing [1, 2]. A key step of the workflow is to use region-specific primers, short single-strand DNA sequences that are by design complementary to a segment in the target region, to capture only the fragments within the target region. However, primer binding and extension can also occur without perfect complementarity, so it is possible for a primer to bind to loci outside the target region where DNA sequences are close or identical to the intended binding site. Off-target templates are amplified and sequenced together with on-target templates, only to generate reads in unwanted regions. Not only do off-target templates waste resources, they take up sequencing capacity that could have been used in the target region, resulting in low read depth and decreased variant calling sensitivity. The degree of freedom in primer design is limited by constraints such as length and GC content, so primers with higher off-target risk are included in some panels more often than others, depending on the repetitiveness of the target region. If we are able to predict the panel-level off-target rate, we can alert scientists of possible off-target outcomes in a quantitative way before they invest time and research fund on high-risk panels. Furthermore, if we are able to predict off-targets for individual primers, we can select the optimal primer(s) among a group of candidates, thus design panels with lower off-target rates.

Off-target rate of a primer depends on the number of possible binding sites and the propensity of hybridization at each site. Hybridization propensity can be predicted using thermodynamic models and characterized by the melting temperature (Tm) of oligos when hybridized to a template. The search of possible binding sites and Tm calculation have been implemented in the widely-used primer design software Primer3 [3]. However, off-target rate is defined by the proportion of reads at unintended binding sites, and read counts cannot be adequately predicted by Tm using a simple linear regression because a portion of information contained in the primer and template sequences is lost when compressed into a single number. We believe an end-to-end approach that predicts the number of reads or unique fragments generated by the hybridization of a primer and a template directly from both sequences would be more likely to succeed. As panel developers with access to a large repository of QIAseq panel sequencing data, we are in a unique position to develop a data-driven, QIAseq-specific off-target prediction model.

Our strategy is to 1) locate possible binding sites for each primer, 2) predict the number of reads at each site, and 3) calculate the primer-level and/or panel-level off-target rate. We developed an in-house algorithm to search for possible binding sites. At each site, we aligned the primer and template sequences and encoded the alignment into an image. We then trained a convolutional neural network (CNN) [4] to predict the read count from the image. Deep learning methods such as CNN are not uncommon in bioinformatics research. They have been used for analyzing how proteins bind to DNA and RNA [5], germline and somatic variant calling [6, 7], off-target prediction of CRISPR-Cas9 gene editing [8, 9], clustering T-cell receptor sequences by antigen [10], and prediction of single-cell DNA methylation states [11]. Our model, named `pordle`, was inspired by some of these works, but the encoding method and network architecture are unique in many ways, as will be explained below. `pordle` was trained using sequencing data on 10 QIAseq panels with 29,274 unique primers in total and then tested in another panel with 7,576 naive primers. The results and methods are presented below.

## 2. Results

### 2.1 Data preparation

We sequenced reference material NA12878 in the 11 panels using the standard QIAseq targeted DNA workflow on Illumina platforms. The reads were processed by the GeneGlobe QIAseq DNA data analysis pipeline [12]. After `BWA-MEM` mapping (GRCh37/hg19), we grouped reads from each primer by the 5’ mapping location (or binding site) and counted the number of unique molecular identifiers (UMI) that were attached during library preparation (Fig. 1a). The UMI counts served as the ground truth in model training and evaluation.

**Figure 1:**
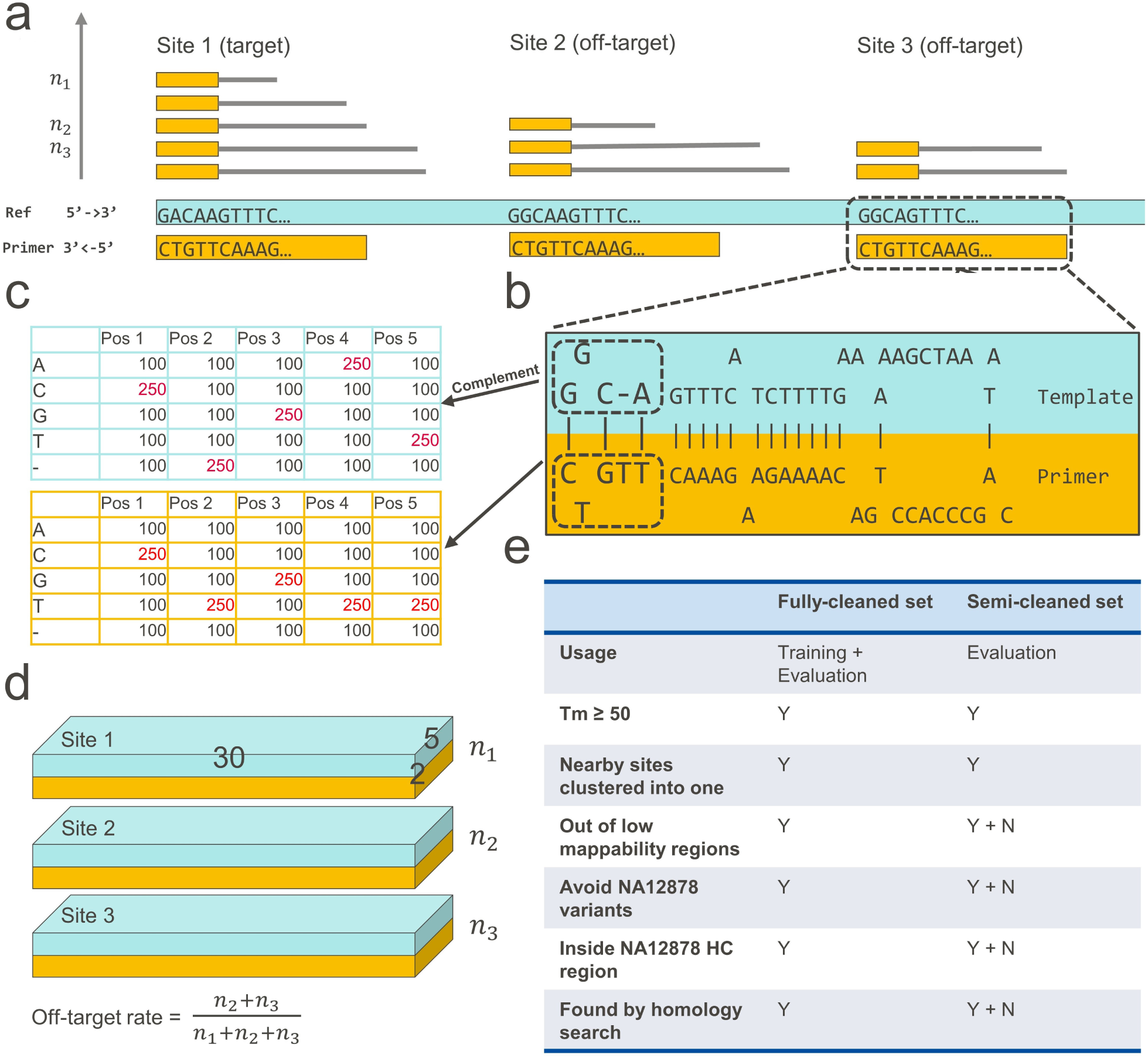
Data processing steps. (**a**) Mapped reads from the same primer are grouped by binding site. The number of UMIs at each site represents primer hybridization propensity and serves as model outcome. (**b**) Primer and template sequences are aligned using `ntthal`. (**c**) The alignment is encoded by two 2D matrices, (**d**) which are stacked into a 3D array that represents a two color channel image as CNN model input. (**e**) A fully-cleaned and a semi-cleaned dataset are generated with different cleaning stringencies for different purposes.

We searched for potential binding sites for all primers in the 11 panels using an in-house homology search algorithm epcr07 (Section 4.1). For the test panel, we ran `epcr07` with two configurations, one being fast and less sensitive (`epcr07-fast`) and the other being slow and more sensitive (`epcr07-slow`). For the training panels, we ran an earlier version of `epcr07` that is close to `epcr07`-slow. Most binding sites found by the homology search are unlikely to be hybridized by the primer due to low Tm, but that is the trade-off for sensitivity. At each site, we found the alignment between primer and template sequences that achieves the highest Tm using `ntthal`, an alignment function in Primer3 based on nearest-neighbor thermodynamical models (Fig. 1b). We believe the `ntthal` alignment is more likely to depict what actually happened in the reaction than the Smith-Waterman alignment and contains crucial information for predicting hybridization propensity. The alignment was encoded into arrays (Fig. 1c) that can be viewed as a multi-channel image (Fig. 1d, Section 4.2), which served as model input.

To compare with image-only models, we combined the image with features that are known or hypothesized to correlate with hybridization propensity, such as the number of mismatches and GC content (Table S1). We scaled the features in training panels to [0, 1] via Minmax normalization and applied the same scaling function to features in the test panel. These features did not add predictive value on top of the image. Cross-validation showed that top-performing models were roughly split 50-50 between image-only and image+feature models and the best model was image-only.

We matched images at possible binding sites to the corresponding UMI counts. For sites without any binding activity (no reads observed), we assigned zero as outcome. We implemented a series of data cleaning steps to the training panels to create a “fully-cleaned” training set where confounding factors were minimized (Fig. 1e and Section 4.3). The fully-cleaned dataset was good for model training and evaluation from a pure analytical perspective. But they are inadequate for assessing the prediction accuracy of a real panel due to considerable amount of noise that were removed from the training process. We created a “semi-cleaned” dataset that, on top of the fully-cleaned set, included sites with low-mappability, overlapping with NA12878 variants, outside of NA12878 high-confidence region, and missed by the homology search (Fig. 1e). The semi-cleaned dataset was still based on the training panels and used together with the fully-cleaned dataset for model evaluation in cross-validation. A fully-cleaned dataset was created for the test panel too.

### 2.2 Model selection by cross-validation

CNN models can be configured by a set of architecture and optimization parameters, each with several options or a range of values (Fig. 2a). Collectively, they can be collectively viewed as the blueprint that guided the construction of each network layer. We will explain the 17 parameters in Section 4.4. We fixed eight parameters via small-scale experiments and left the other nine to be determined by cross-validation. Instead of exhausting all 5,760 parameter combinations, we randomly selected 47 to enter the cross-validation and chose the one with the best validation performance as the final model. Individually, most parameters had insignificant or inconsistent impact on prediction accuracy in fully-cleaned and semi-cleaned held-out sets (Fig. S1). The only notable observation is that mean square error (MSE) as loss function reduced panel-level prediction error by 50% compared to logcosh.

**Figure 2:**
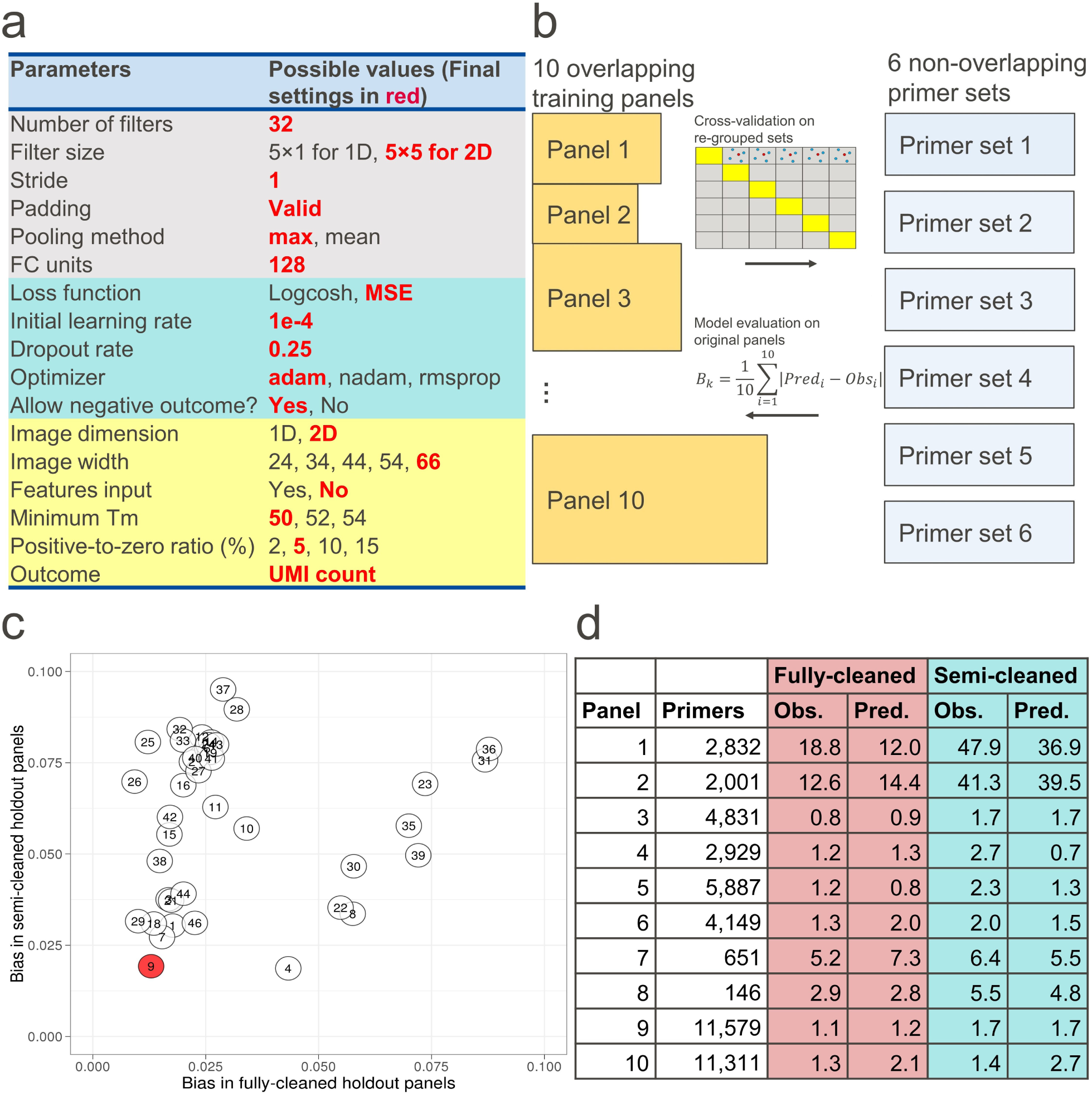
Model selection by cross-validation. (**a**) Model parameters and values. Parameters with more than one values are determined by cross-validation. Gray: architecture parameters. Green: optimization parameters. Yellow: data input parameters. (**b**) 10 training panels were re-grouped into 6 non-overlapping sets of primers for a 6-fold cross-validation. After completion, models were evaluated by the absolute prediction bias of off-target rates, averaged across all 10 *original* panels. (**c**) Parameter set (#9) achieved the best overall performance in fully-cleaned and semi-cleaned held-out sets and was chosen as the final setting. (**d**) Panel-level off-target rate prediction of the final ensemble model.

Because some primers belong to multiple panels, we regrouped the primers in the 10 training panels to 6 non-overlapping, similar-sized sets by dividing larger panels and combining smaller ones. We performed a 6-fold cross-validation that iteratively held out one primer set for evaluation and the other five for training (Fig. 2b). Normally in cross-validations, models are evaluated by the same training loss function. In our case, however, the model with minimum MSE tended to underestimate the panel-level off-target rate due to overwhelming zero and low UMI sites. So we used panel-level prediction bias to assess model performance.

Upon the completion of the cross-validation, every site had been predicted by a model trained elsewhere, so we calculated the predicted off-target rate for each *original panel* from site-level UMI predictions. We used the mean absolute prediction bias (MAPB) across all 10 training panels for model selection. The optimal parameter set (#9) achieved a MABP of 1.3% in fully-cleaned held-out sets and 1.9% in semi-cleaned sets (Fig. 2c). Under this setting, the models were able to predict high-risk panels (1, 2), mid-risk panel (7), and low-risk panels (3-6, 8-10) with reasonable accuracy (Fig. 2d). Some reports suggested training a final model using all data [5], but we found that such a model performed poorly in the test panel. A possible reason is that due to the lack of a validation set for loss monitoring, the training did not stop at the right timing, which resulted in under-fitting or over-fitting. We adopted an ensemble approach and took the median of the six model predictions as the final prediction.

### 2.3 Prediction accuracy on test panel

The test panel had an overall off-target rate of 10.6% and `pordle` predicted 10.5% with the fast configuration of `epcr07` and 11.4% with the slow configuration of `epcr07` (Fig. 3). If we focus on a fully-cleaned subset of binding sites, the observed off-target rate was 4.3% with fast `epcr07` and 4.6% with slow `epcr07`. The two numbers differ because sites in the fully-cleaned dataset must be found by the homology search, and the slow configuration found more sites. The prediction for the fully-cleaned dataset was 4.0% with `pordle-fast` and 4.8% with `pordle-slow`. We also trained support vector regression (SVR) models with linear and radial basis function (rbf) kernels using the same training data (Section 4.7). The SVR model with linear kernel predicted 51.9% off-target rate for the fully-cleaned dataset and 61.1% for the original dataset. SVR with rbf kernel gave even higher predictions. While `pordle` is expected to outperform traditional machine learning methods like SVR, the performance of SVR may be under-estimated because the data cleaning parameters (minimum Tm = 50 and positive-to-zero ratio = 5%) that worked for CNN may not be optimal for SVR.

**Figure 3:**
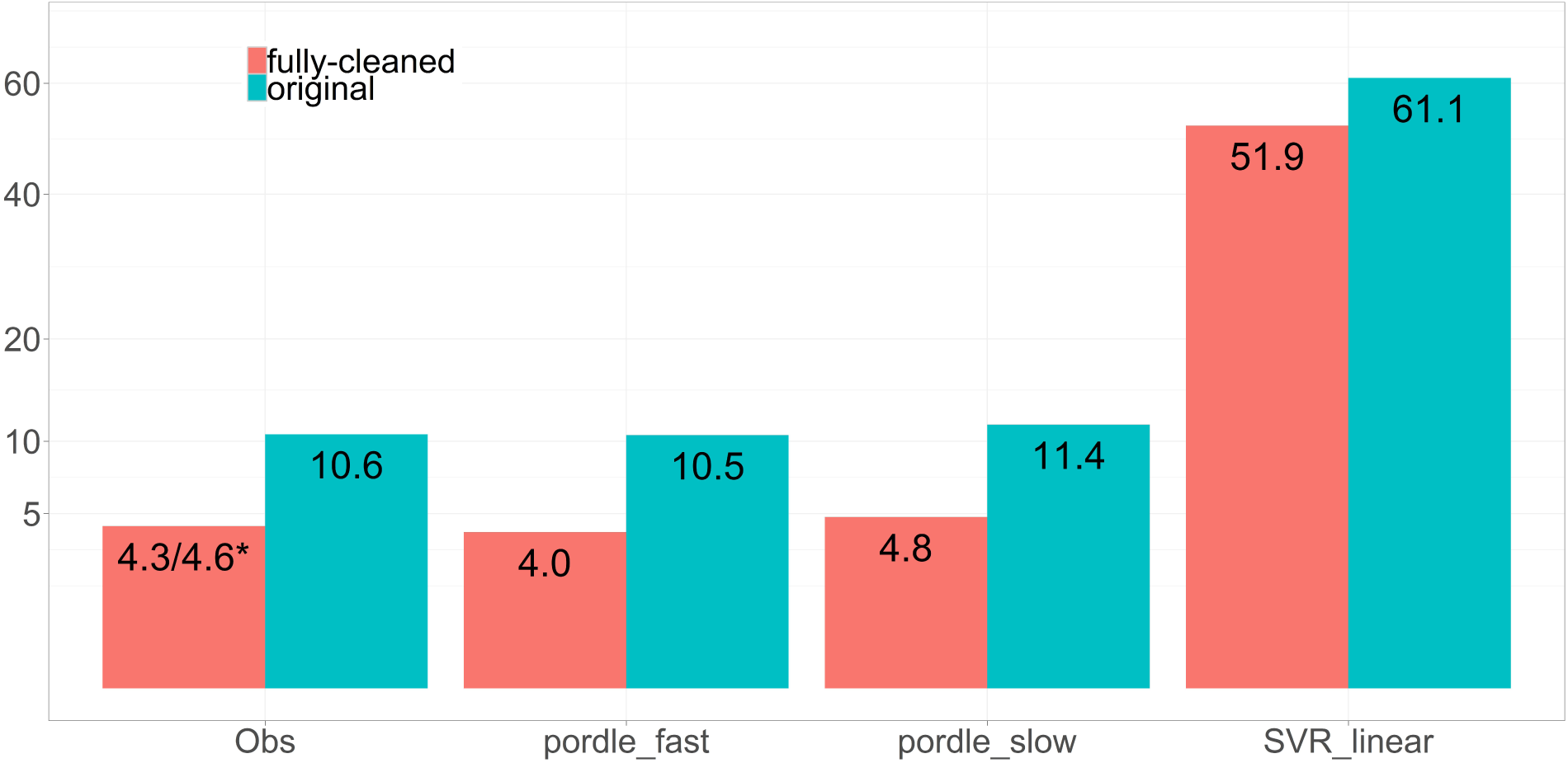
Panel level prediction results. The observed off-target rate in the fully-cleaned test panel was 4.3% with the fast configuration of `epcr07` and 4.6% with the slow configuration. SVR with rbf kernel produced even higher predictions than linear kernel, hence not shown here.

The low prediction bias by `pordle` should be interpreted with caution. If we restrict to the sites with Tm *≥* 50 and found by `epcr07-fast` (without further cleaning), the true off-target rate dropped from 10.5% to 7.2% (7.6% for `epcr07-slow`). The 3.3% discrepancy was about equally contributed by the Tm filtering and being missed by `epcr07`. Therefore, if `epcr07` was configured to be more sensitive and located all binding sites, `pordle` would have predicted a higher off-target rate.

We investigated how often can `pordle` pick from a pair of nearby primers the one with lower off-target rate. By our definition, “Nearby primers” are within 250bp if on the same strand and within 500bp if on opposite strands. We collected all pairs of nearby primers in the test panel whose true off-target rates differ by above some threshold, predicted the off-target rate for each primer, selected the primer in each pair with lower predicted off-target rate, and counted the number of pairs the selected primer’s true off-target rate was indeed lower than the other primer. For nearby primers that differ by at least 1%, `pordle` had over 80% chance selecting the better one. The better primer was selected in 89.2% of 3,835 pairs of nearby primers whose off-target rates differ by at least 10% (Fig. 4a). The rank-preserving percentage increased along with the gap of true off-target rate until the gap reaches 60%, when it started decreasing slowly. The rank-preserving percentage tumbled to 82% for the 201 pairs of primers with gap *>* 80%, indicating poor prediction for some high off-target primers. This pattern was not observed in the fully-cleaned data, in which the rank-preserving percentage kept increasing as the gap increases.

**Figure 4:**
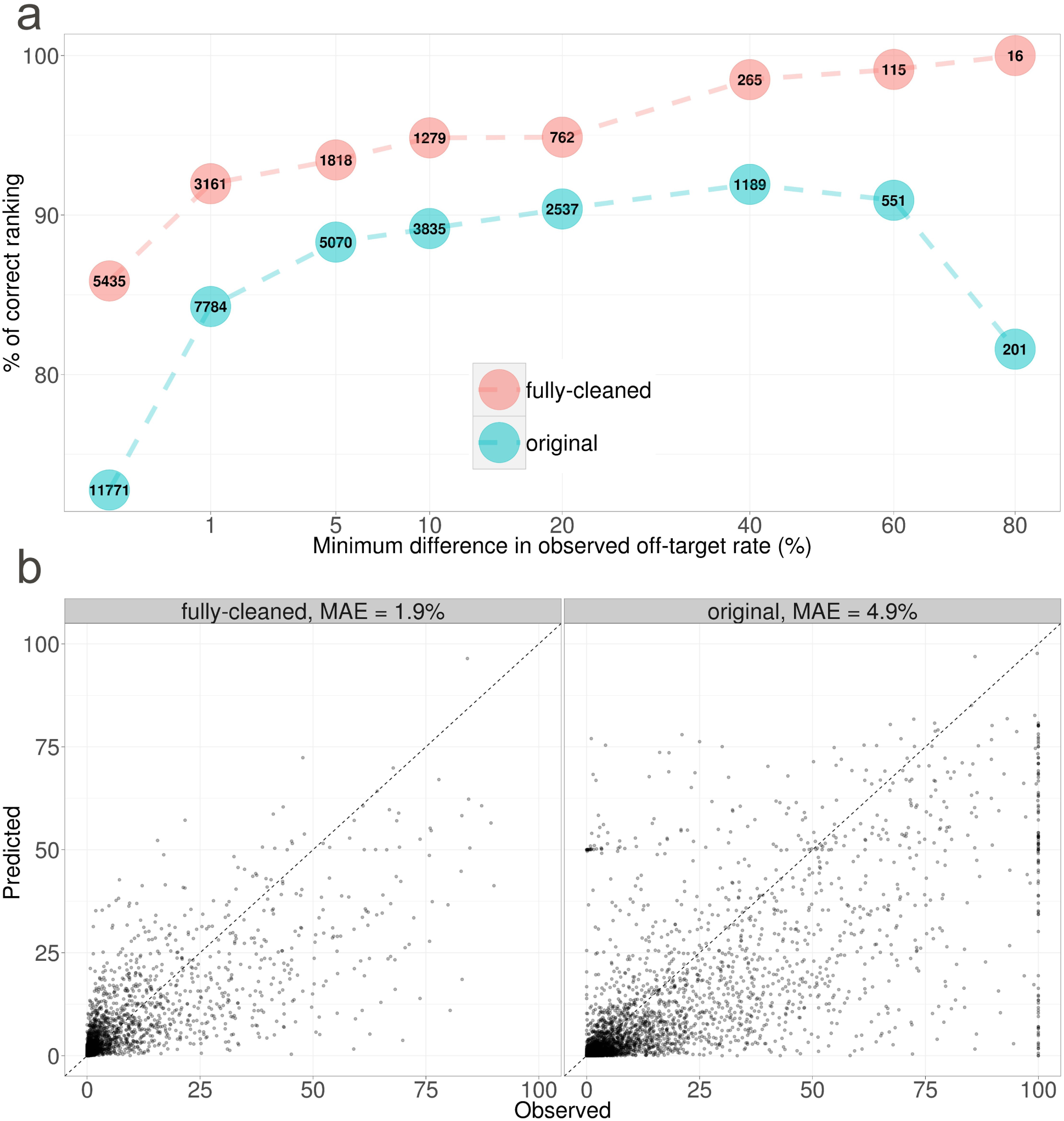
Primer level prediction results. (**a**) The figure answers the question: given a pair of nearby primers (within 250bp if on the same strand and 500bp if on opposite strands) in the test panel, how often can `pordle` correctly pick the one with lower off-target rate. In each circle is the number of nearby primer pairs whose observed off-target rates differ by at least the corresponding x-axis value. The corresponding y-axis is the percentage of such primer pairs for which `pordle` correctly predicted the order of off-target rates. (**b**) Predicted versus observed off-target rate (%) for each primer in the test panel. MAE stands for mean absolute error, which is the absolute value of prediction bias averaged across all primers.

The primer-level prediction was more accurate in the fully-cleaned data with a mean absolute error of (MAE) 1.9% (Fig. 4b). In the original data, MAE increased to 4.9% and the model under-predicted many high off-target primers. Notably, some primers with off-target rate = 1 were severely under-predicted.

The predictions were less reliable for primers with low mappability at the intended binding site (Fig. 5, top panel). Mappability was measured by a composite CRG40 score between 0 and 1 (UCSC Genome Browser CRG track, smaller values indicate less mappability of 40mers). We grouped the absolute bias into three levels: low (0-10%), medium (10-50%) and high (50-100%). The majority (82.5%) of low bias primers had CRG40 = 1, while more primers in the medium and high bias groups had CRG40 below 0.5. Low mappability at the intended binding site could inflate the off-target rate because 1) some on-target reads may be mapped to other loci and 2) the correctly-mapped on-target reads may have MAPQ*<* 60 due to non-unique mapping loci and discarded in our analysis. We did observe higher spikes of low MAPQ reads at the intended sites for medium and high bias primers (Fig. 5, bottom panel).

**Figure 5:**
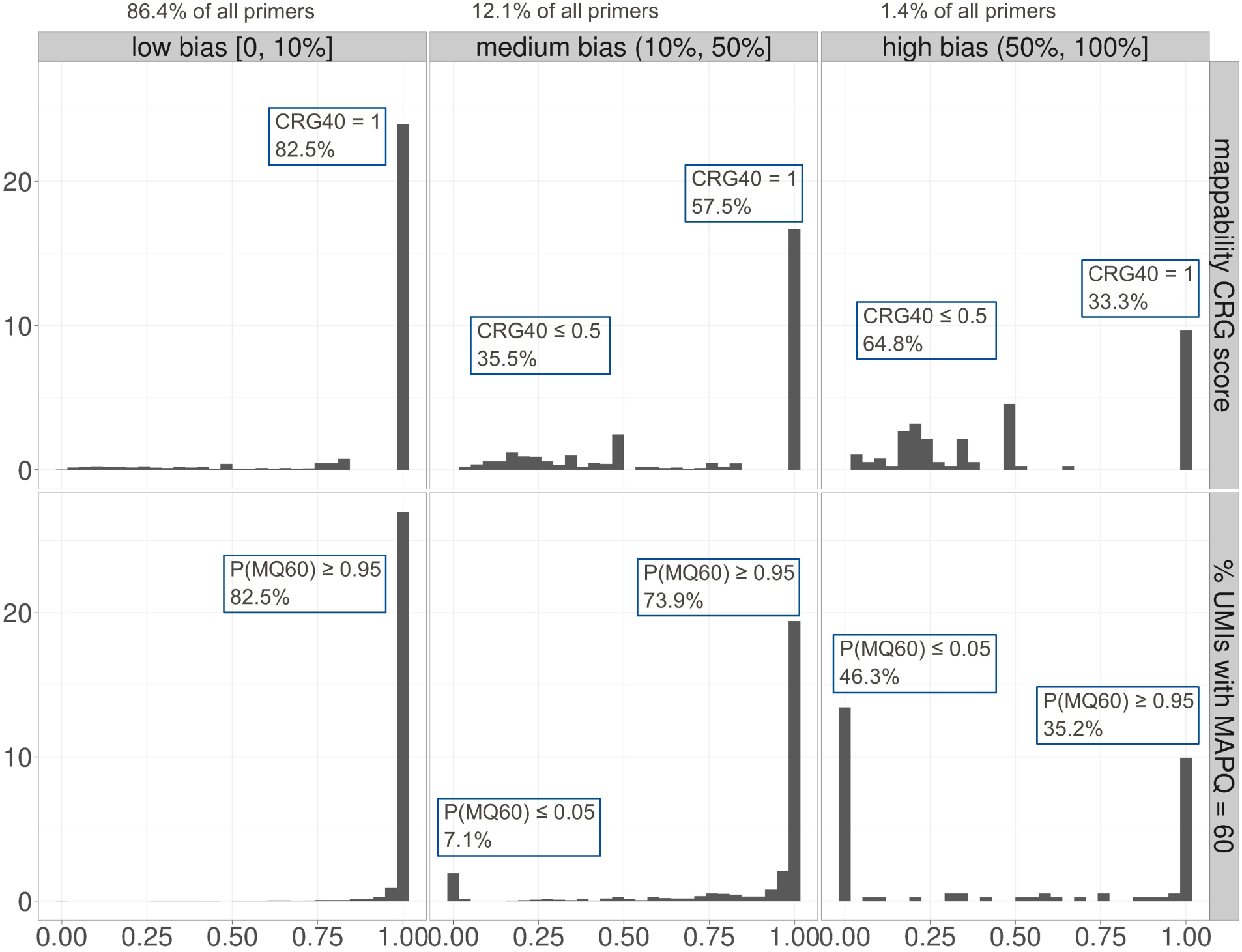
Primers with low mappability (measured by a composite CRG40 score) and/or a high percentage of low mapping quality reads at the intended binding site. The histograms and data are based on the original test panel data.

In other words, some of the high off-target primers actually had lower off-target rates if low MAPQ reads were counted in. This partially explained the sudden drop of correct ranking percent at gap = 80% in Fig. 4a and the under-prediction of primers with off-target rate = 1 in Fig. 4b. This pattern was less often seen in the fully-cleaned data because many of such primers were excluded.

## 3. Discussion

We developed pordle, a deep learning model that predicts the off-target rate for QIAseq targeted DNA sequencing panel or individual primers. In an independent test panel with 7,576 primers, `pordle` accurately predicted the panel level off-target rates with a bias of -0.1%. It also showed good accuracy in predicting low off-target rate primers and had the ability to preserve the ranking of two nearby primers. We believe `pordle` can add value to the QIAseq DNA panels and its quantitative prediction for individual primers can help internal panel design work.

The results should be interpreted with caution due to two complications in the methodology and data processing steps. First, `pordle` relies on the homology search algorithm to find nearly all binding sites. A portion of observed off-target sites in the test panel were not found by `epcr07`, making the panel-level prediction look better than it actually is. Had more binding sites were found, the prediction bias would have been higher. Second, because reads with MAPQ ¡ 60 were discarded to keep only perfectly-mapped reads in the analysis, off-target rates for some primers with low mappability at the intended binding site were counted too high. Therefore, the prediction for some high off-target primers are better than what appeared in Fig. 4b. The primer-level prediction was much more accurate in the fully-cleaned dataset, implying that a major source of bias came from the “noisy” sites and primers (Fig. 1e) that were not included in the training.

As a deep learning-based model, `pordle` takes the image converted from the primer-template alignment as the only input without relying on any manually curated features. The image-only model outperformed image+feature model in cross-validation and greatly outperformed the feature-only SVR model in the test panel. This result is not surprising because deep learning models have been shown to outperform traditional machine learning models in applications where the information contained in the complex data cannot be fully captured by manually selected features. In our case, the thermodynamic features Δ*G*, Δ*H*, Δ*S* added new information to the alignment because they were calculated using formulae that were derived by general physical laws. But such subtle additions did not lead to better performance in the cross-validation.

The CNN architecture (Fig. 6) was determined by trial and error in a rather ad-hoc manner. We are convinced that the convolution layers are crucial to the learning because networks with only dense layers did not achieve the same level of prediction accuracy. Deeper networks with as many as 8 convolution layers did not improve the prediction either. This is possibly because our dataset is not big enough and the image converted from the alignment is not complex enough to fully exploit the power of deeper networks. Nevertheless, given more time and resources, we could explore more modern, complex network architectures such as ResNet [13] and/or take advantage of existing networks using transfer learning.

**Figure 6:**
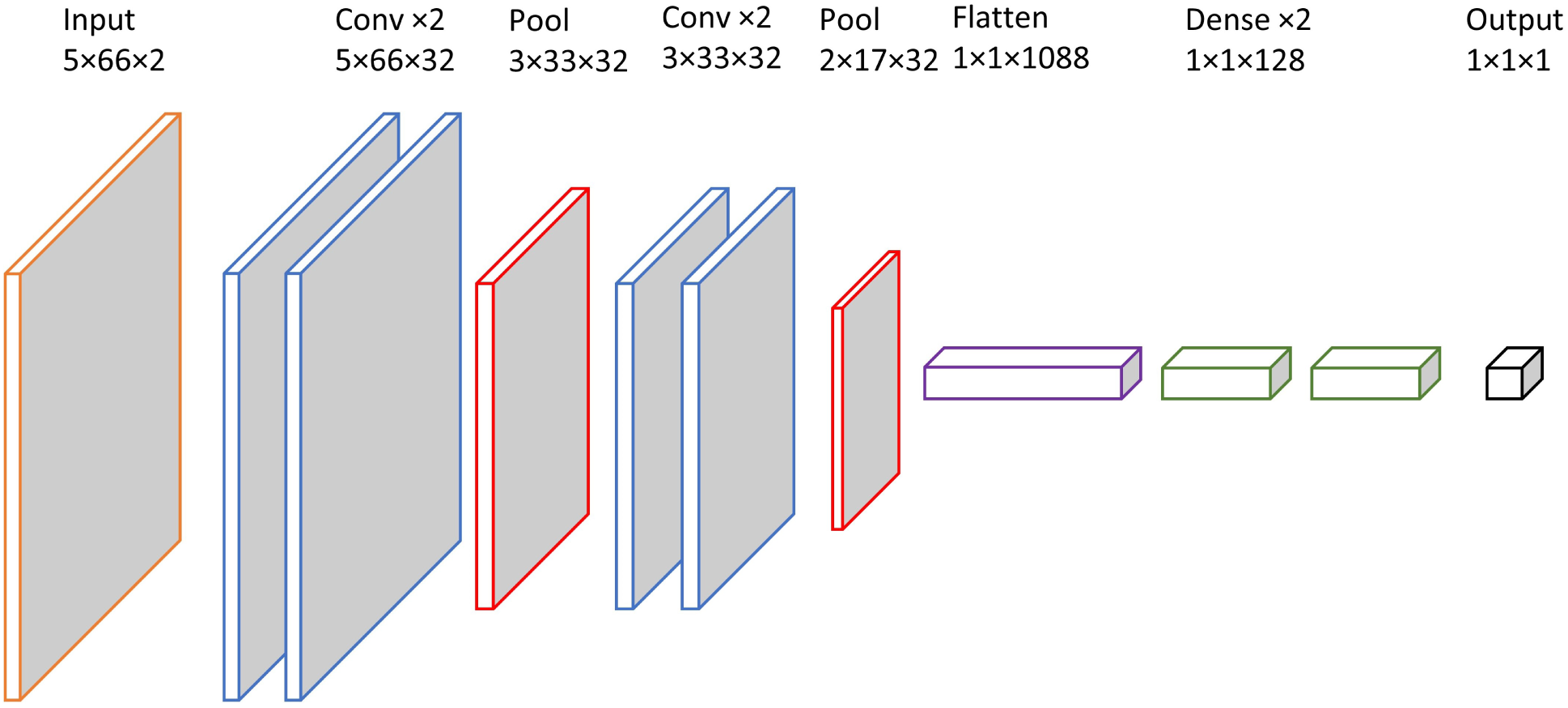
CNN model architecture for the final setting.

In the future, we can test `pordle` on more datasets to better assess its performance. We can also join additional datasets with the current 11 panels to re-train `pordle` on a bigger training set. The image encoding method and model architecture described here can potentially be applied to the prediction of gRNA off-target in CRISPR-Cas9. The two problems are similar in that both require the prediction of reads/UMIs based on the mismatch pattern between two sequences.

## 4. Methods

### 4.1 Homology search algorithm epcr07

We used epcr07, an in-house homology search algorithm to find potential priming sites. The algorithm implements a simple “seed and extend” method in 4 steps. First, we design seed mismatch patterns that exclude 3’ side mismatch patterns rarely observed, presumably due to lack of Taq enzyme recognition. This seed strategy provides high search sensitivity, but does not generate primer-template sites that are very unlikely to generate PCR product. Second, we extract genome sequence at seed site. Third, we drop primer-template pairs that are very unlikely to be thermodynamically stable. Finally, we estimate primer-template thermodynamic stability. For primer-template hits passing the previous step, we perform a nearest-neighbor thermodynamic stability calculation and output genome hit sites above the desired stability threshold.

We ran two configurations of `epcr07`, one with the SSW specificity filter disabled and the other with it enabled. If the SSW specificity filter is disabled, `epcr07` runs much slower but finds more sites.

### 4.2 Image encoding

To apply CNN, we converted the alignment into an array, or image (Fig. 1c). The conversion method is similar to a one-hot encoding, where each alignment position is coded by a 5-bit vector corresponding to four nucleotide acids (ACGT) and deletion (-). The correct category takes value 250 and others take value 100. The primer and template were converted to 2D matrices separately before stacking together, resulting in a 3D array that represents an image with two color channels (Fig. 1d). To make equal-sized images as required by CNN, we padded zeros to the right of shorter primers and/or trimmed bases from the less important 5’ end (right side in Fig. 1b) of longer primers. We also padded 4 bases of zeros to the left of all images for the sliding filter to learn fine features near 3’ end of the primer/template alignment. Finally, we divided scaled the input array between [0, 1] by dividing 255 everywhere.

Alternative to 2D encoding (Fig. 1c, d), we could convert the `ntthal` alignment to an 1D image with 3 channels. The first two channels represent the primer and template sequences, where the four nucleic acids A, C, G, T are coded by 250, 180, 100 and 30. The third channel encodes the type of match at each position. Match (or complementary), mismatch, insertion and deletion (template as reference) are coded by 250, 180, 100, 30. The 1D image has the same length as the primer before padding or trimming to a uniform length. The CNN architecture for 1D input is similar as 2D except for 5 × 1 kernels instead of 5 × 5.

### 4.3 Data cleaning steps

First, we adjusted the ratio between positive- and zero-UMI sites by excluding any site below Tm 50 °C, followed by further downsampling zero-UMI sites until certain positive-to-zero ratio is achieved (2, 5, 10, 15%). Real binding sites below Tm 50 were removed as well, but we sped up the training process and more importantly, avoided a falsely accurate predictive model that predicts near-zero for every site. Further downsampling of zero UMI sites risks putting too heavy weights on high UMI sites. After trial-and-error, we settled with a moderate positive-to-zero ratio. Second, we excluded sites where the primer/template alignment may not be adequate for prediction, including those overlapping with known NA12878 variants, outside the high-confidence regions (GIAB v3.3.2), and in low mappability regions (characterized by CRG score [14], Methods). We also dropped primers whose on-target UMIs are below 10% of the median across all primers (normalized across panels) as outliers. Third, we merged off-target sites within 15 bases from the target sites and counted their UMIs as on-target, because many of such reads were actually on-target but mapped a few bases away due to nearby short repeats. Similarly, we combined off-target sites within any 10-base interval and summed up their UMIs. Fourth, we kept only reads with perfect mapping quality (60) to prevent false off-target UMI count due to mapping errors. We normalized UMI counts in each panel to the same median on-target UMI. After normalization, we took the mean UMI at each site for primers in multiple panels.

### 4.4 Parameter space for model selection

Throughout the study, we considered six architecture parameters including the number, size, and moving stride of filters, padding and pooling method and the number of units in fully connected (FC) layers. The filter parameters determine how local features are learned in the convolution step. Padding method determines how filters behave on the edge of the image. Pooling method determines how to extract information and reduce dimension in the pooling layer. The architecture parameters are further explained in Section 4.5. We considered five optimization parameters: loss function, optimization algorithm (optimizer), initial learning rate, dropout rate and whether to allow negative prediction outcome. Loss function defined a quantity of prediction error to be optimized and others determined different aspects of the optimization algorithm. Unique to this problem, we had six additional parameters regarding the input data format: image dimension, image width, whether to include artificial features as input, minimum Tm, downsampling rate and outcome quantity. The first five parameters were explained in Section 4.3. The last one, outcome quantity, could be either the absolute UMI count (the chosen one) or the relative UMI count as a ratio to the intended binding site of the primer.

### 4.5 CNN architecture

After exploring several options, we settled on a basic CNN architecture that consists of four convolution layers, two pooling layers and two fully connected layers (Fig. 6). The final model 2D images, in the first convolution layer, thirty-two 5 × 5 × 2 filters moved across both channels of the image one step a time (stride = 1) and convolved with every 5 × 5 × 2 cube in the image (dot product,∑_*i,j,k*_ *x*_*ijk*_*y*_*ijk*_) plus an intercept to generate a stacked feature map. Zeros were padded to the edges of the input image so that the feature maps had the same height and width as the input. This layer had 32 × (5 × 5 × 2 + 1) = 1, 632 parameters and output via ReLu activation (*x*^+^ = max(0, *x*)) a stacked feature map of size 5 × 66 × 32. The purpose of the convolution layer was to extract features from short sequences (say 5-mers) that associate with binding propensity. The feature map was fed into a second convolution layer to learn more abstract features. This layer had 32 5 × 5 × 32 filters with 32 × (5 × 5 × 32 + 1) = 25, 632 parameters and output a stacked feature map of the same size as input. A pooling layer followed after two convolution layers to downsize the feature map by taking the maximum or mean of non-overlapping (stride = 2) 2 × 2 squares in each slice of the map. A 66-bit vector of zeros was padded to one of the long edges, making the output of the pooling layer a 3 × 33 × 32 array. We stacked up another round of two convolution and one pooling layers. These two convolution layers had 32 3 × 5 × 32 filters, each bringing in 15,392 parameters. A dropout layer that randomly set 25% of the pixels in the feature map to zero was inserted after each round. Applying dropout on convolution layers is less common than on fully connected layers due to lack of clear interpretation, but it has been reported to help reduce prediction error [15]. The output of the second pooling layer, a 2 × 17 × 32 array was flattened into a 1D vector and fed into two FC layers with 128 units each. The first FC layer introduced (2 × 17 × 32 + 1) × 128 = 139, 392 parameters and the second 129 × 128 = 16, 512 parameters. If artificial features were used for prediction (Fig. 2a), they were concatenated with the flattened convolution output before entering the FC layers. Finally, the output of the entire network was calculated by a linear combination of the 128 neurons (129 parameters) from the second FC layer with identity activation. The whole network under the final setting had 214, 081 parameters (weights) for optimization.

### 4.6 CNN model training

In each training session, we allocated 80% of the sites (“training set”) for model training using a stochastic gradient descent algorithm that aims to optimize a loss function, which is some form of prediction error plus an optional penalty on model complexity. During the training, we evaluated model performance with the other 20% sites (“validation set”, not to be confused with the held-out primer set or the test panel) by calculating the loss after every epoch. When validation loss stagnates, we can stop training to avoid over-fitting or reduce the learning rate (step size of gradient descent) to further find tune the weights (Fig. 7). We set the training to stop after validation loss stayed plateau (non-decreasing) for 10 consecutive epochs and took the weights at 10 epochs prior to termination as final weights. We also set to decrease the learning rate by a factor of 10 each time validation loss plateaus for 5 epochs.

**Figure 7:**
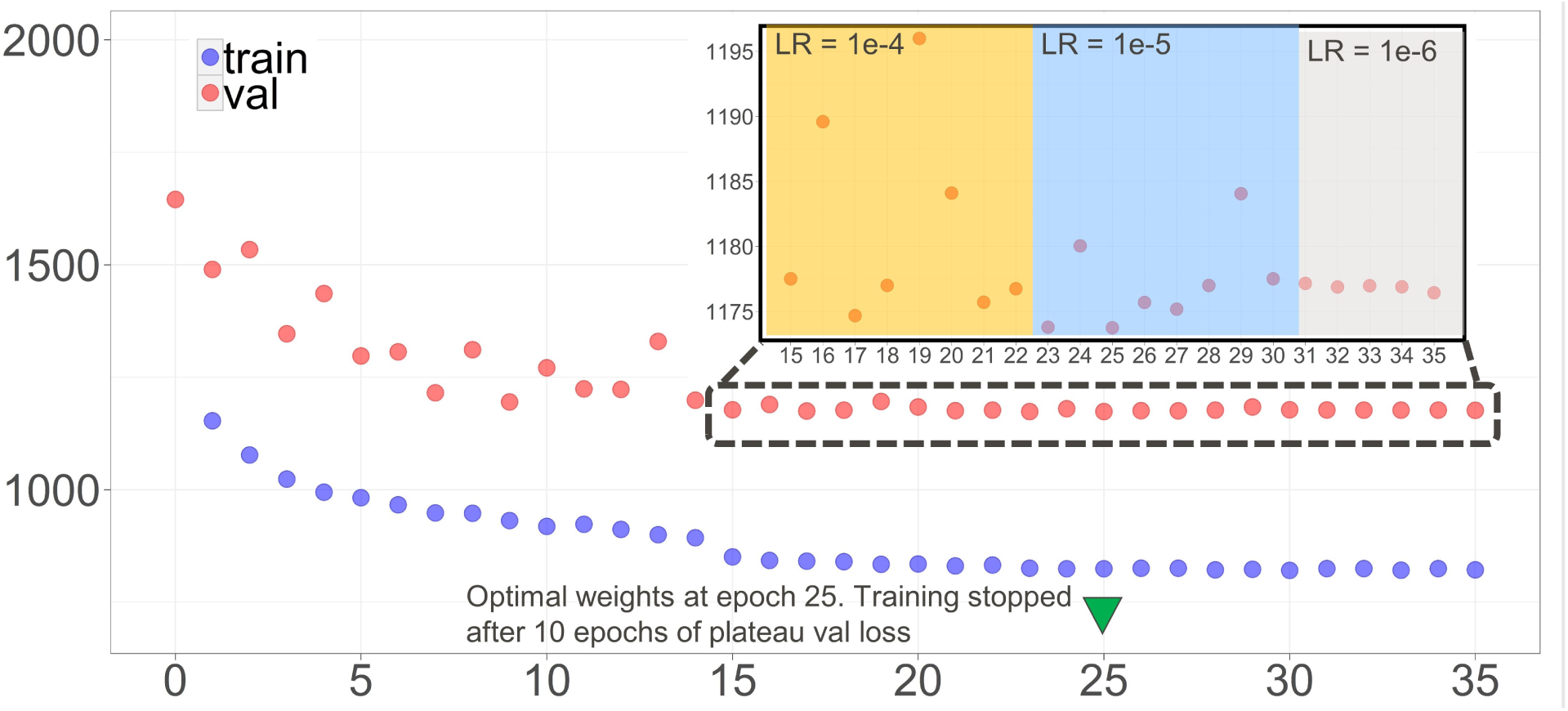
An example of a training session. Training stops after validation loss plateaus (non-decrease) for 10 epochs. Learning rate decreases 10-fold after validation loss plateaus for 5 epochs.

We used Keras with an R interface and TensorFlow backend for CNN model training. All computations were carried out on CPUs of a 64-core workstation. Training time varied by model complexity and data size, which were widely different among the 47 parameter sets. The model with the optimal parameter set (Fig. 2a) took 26-57 minutes to train using all 64 cores. According to our tests on the Google Cloud Platform, training time could be reduced by about 60% with a single Nvidia Tesla T4 GPU even though the GPU usage was usually below 30%.

### 4.7 Support vector regression

We trained a support vector regression (SVR) model using the training panels and applied it to the test panel. The SVR model takes 22 manually curated features (Table S1) as input to predict the UMI count at a site. We used the same dataset as in the optimal CNN parameter setting (Fig. 2a) to train the SVR model, acknowledging that the data processing parameters (minimum Tm = 50 and positive-to-zero ratio = 5%) that worked for CNN may not be optimal for SVR. We did not perform a similar scaled cross-validation study for SVR because with over half a million data points, training an SVR model with radial basis function (rbf) kernel is very time-consuming (26 hours for one model using 24 cores, after 30 hours for a grid search for optimal model parameters).

If we denote the feature vector at site *i* as *x*_*i*_ and the UMI count as *y*_*i*_, the SVR model aims to find a set of weights *w* (22-bit vector) and intercept *b* that minimizes the loss function

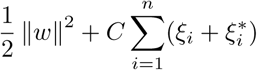

subject to constraints that for each *i*,

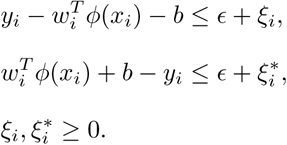

In the above, 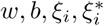 are to be optimized and *C, ∊, φ*(·) are fixed parameters. *∊* > 0 is the desired maximum distance between the prediction 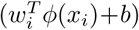 and the true value (*y*_*i*_). *C* > 0 determines the trade-off between model complexity (*∥w∥*^2^) and the total amount of prediction bias that exceeds *∊* (soft margin), represented by the sum of slack variables 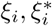. *φ*(·) is a function that maps *x*_*i*_ to a higher dimensional space and *K*(*x*_*i*_, *x*_*j*_) = *φ*(*x*_*i*_)^*T*^ *φ*(*x*_*j*_) is the kernel function.

We chose two commonly used kernel functions: linear kernel 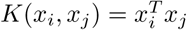 and rbf kernel *K*(*x*_*i*_, *x*_*j*_) = exp(*-γ*∥*x*_*i*_ *-x*_*j*_∥^2^. After min-max scaling the feature vector to [0, 1], we performed a grid search with a fold cross-validation to find the optimal parameters that minimizes the mean square prediction error. The grid search ranges were (0.1, 1, 5, 10, 25, 50, 100) for *ϵ*, (0.05, 0.1, 0.25, 0.5, 1.0) for *γ*, (100, 1, 000, 2, 500, 5, 000, 7, 500, 10, 000) for *C* with linear kernel and (10, 000, 25, 000, 50, 000, 100, 000) for *C* with rbf kernel. The final models, SVR-linear and SVR-rbf, were trained using the optimal parameters on the whole training data. The training of linear kernel SVR was relatively fast and completed using Python module `scikit-learn`. For the rbf kernel SVR, we used `scikit-learn` for grid search and C++ library LIBSVM for faster model training.

## Supplementary Tables and Figures

**Table S1.**
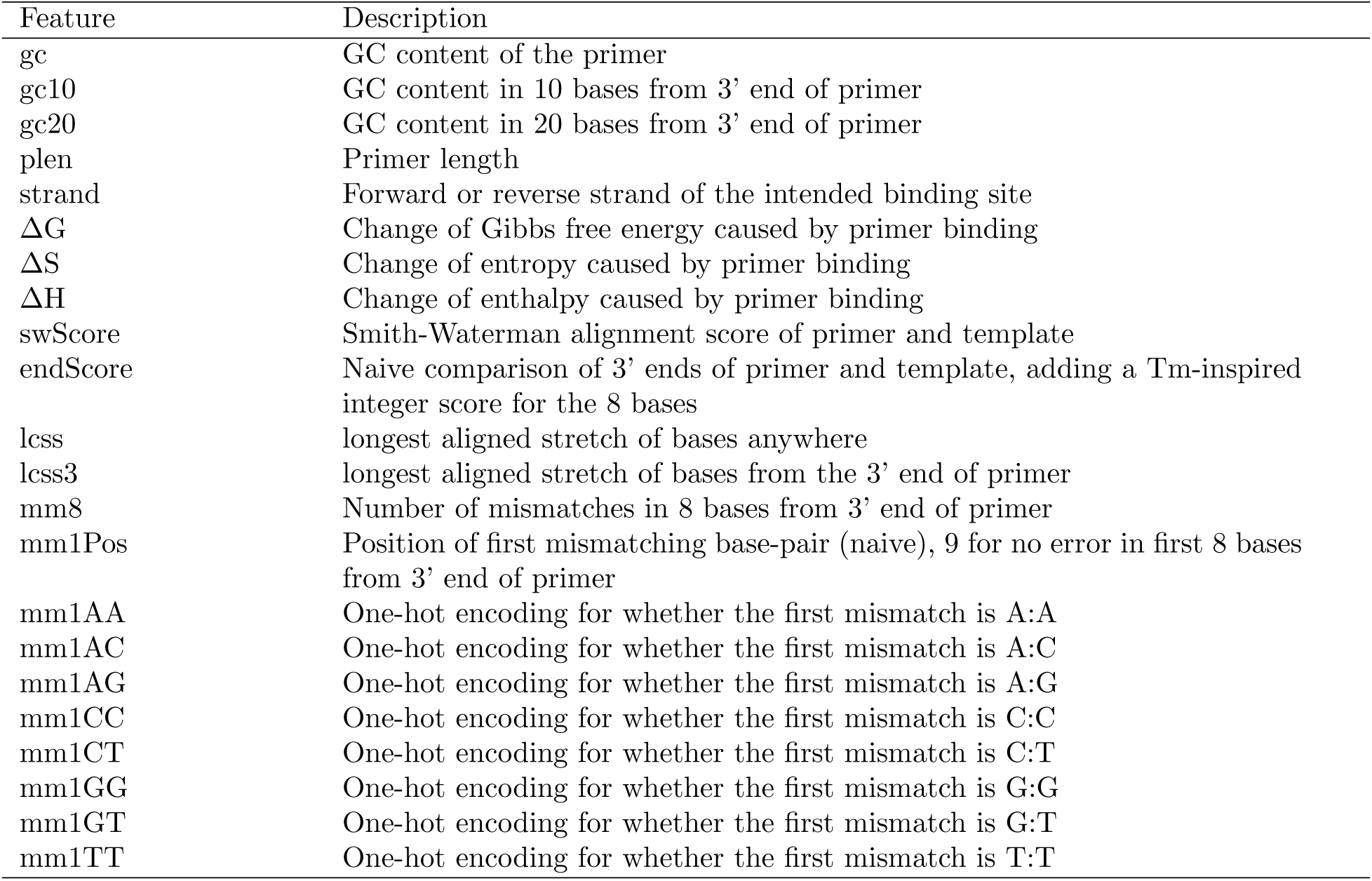
Manually curated features for joint modeling

**Figure S1:**
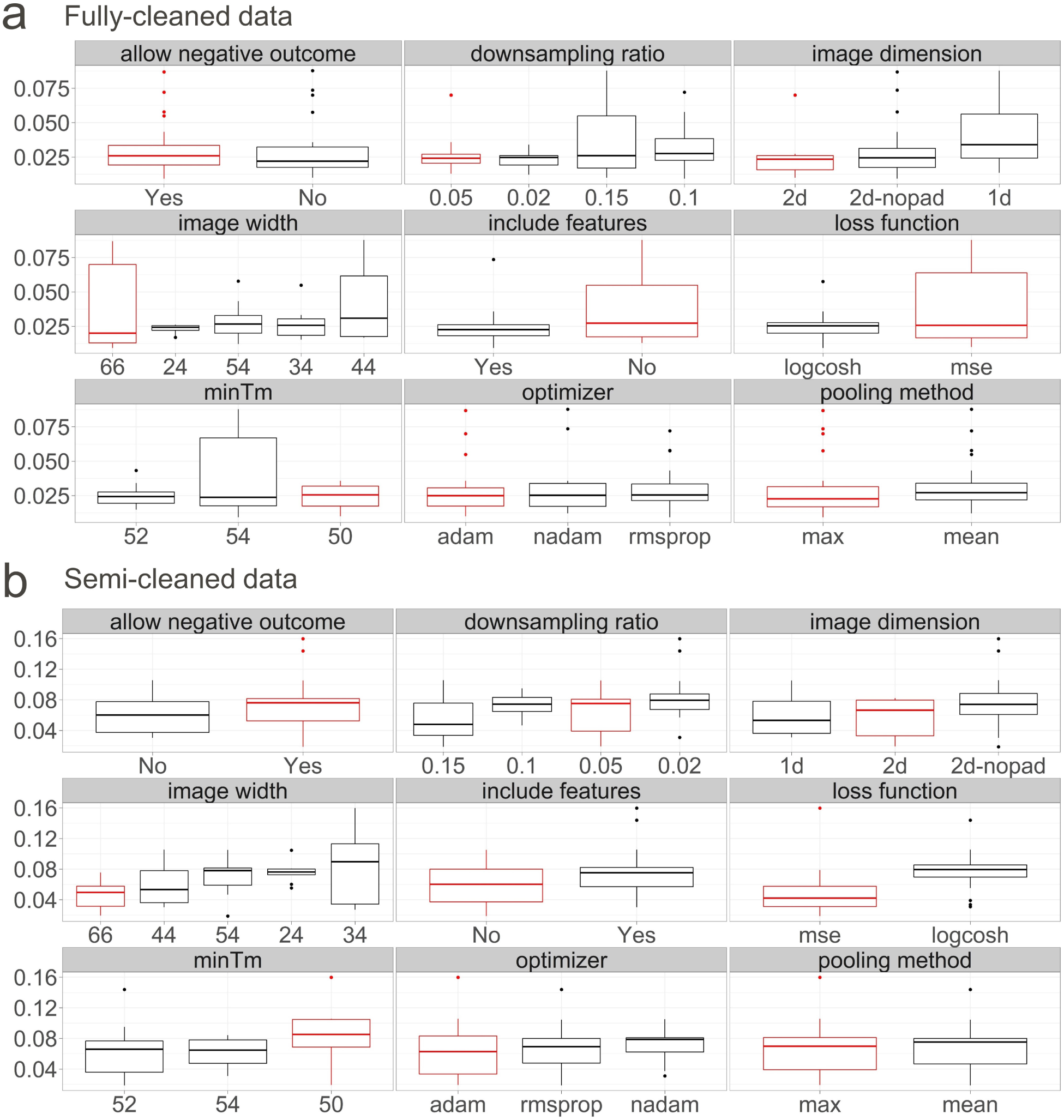
Mean panel-level absolute prediction bias by individual parameter for (**a**) fully-cleaned and (**b**) semi-cleaned held-out primer sets. An outlier with mean absolute prediction bias above 0.3 was not shown in (**b**). The final parameter values were shown in red.

